## Supplementary figures and images for "Metrics of High Cofluctuation and Entropy to Describe Control of Cardiac Function in the Stellate Ganglion"

### Supplemental File 1

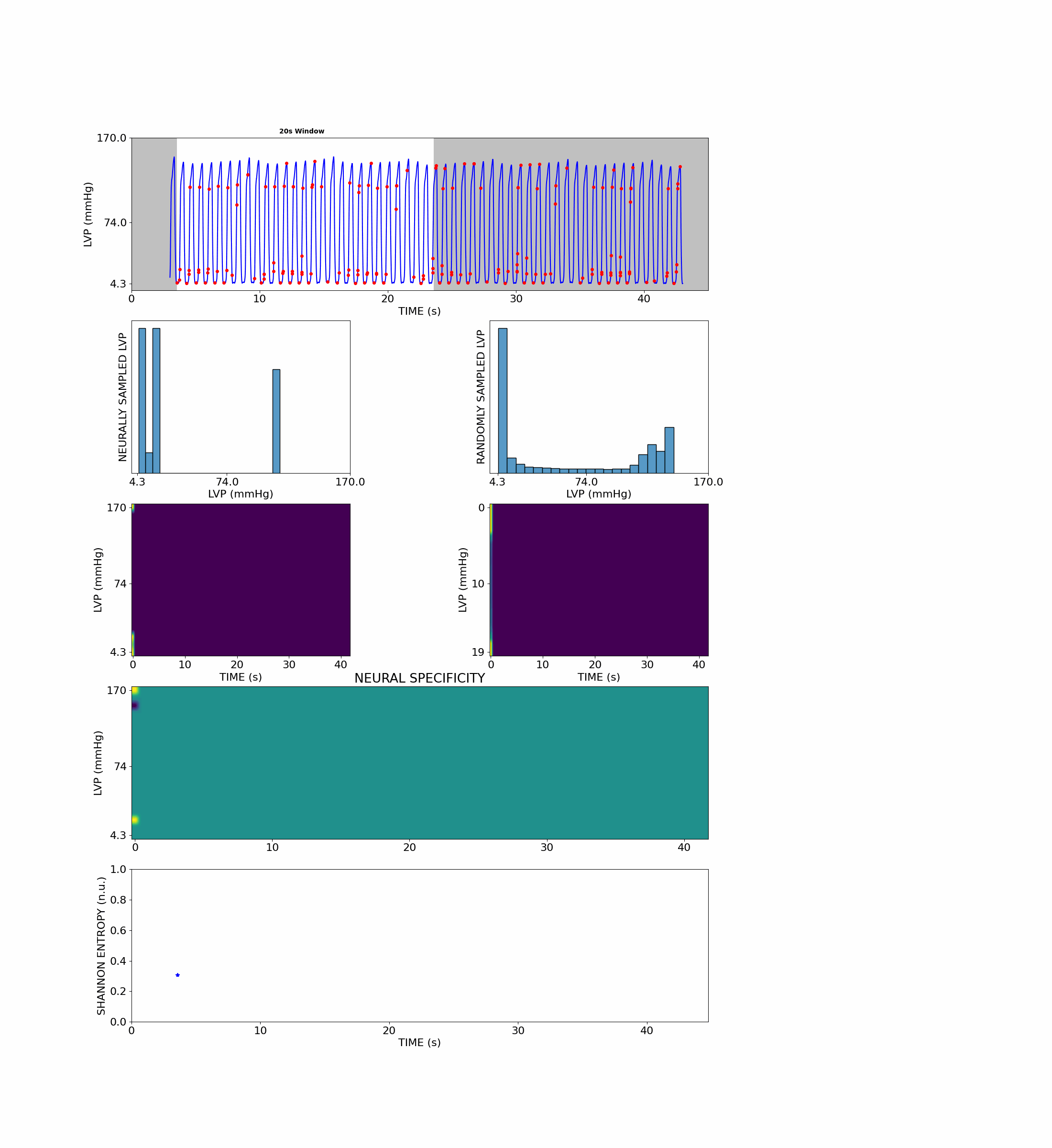
